# Deep learning-based behavioral analysis in a neonatal rat model of hypoxic-ischemic brain injury

**DOI:** 10.64898/2026.04.07.716979

**Authors:** Benjamin Lee, Hang Xing, Bingqing Wang, Matthew Lam, Xiaodi F. Chen

**Author notes:** Authors contributed equally to this work (Benjamin Lee, Hang Xing). **Corresponding Author:** Xiaodi F. Chen, MD, PhD.

## Abstract

Hypoxic-ischemic (HI) brain injury in neonates is one of <u>the</u> leading causes of lifelong neurological disability. Behavioral tests in preclinical rodent models are widely used to assess motor and cognitive outcomes after HI injury; however, these assays usually depend on subjective and labor-intensive manual scoring. Recent advances in markerless pose estimation offer new opportunities for automated and reproducible behavioral quantification in animal and infant recordings, but their use in neonatal HI preclinical studies remains limited. Wistar rat pups underwent HI injury using the Rice-Vannucci model at postnatal day 7 (P7). Three developmental behavioral tests included righting reflex (P8), negative geotaxis (P14), and wire hang (P16), were recorded and analyzed by both a human rater and an automated pipeline using DeepLabCut (DLC), an open source markerless pose estimation framework. Automated measurements were compared with manual scores using Intraclass Correlation Coefficients (ICC), Bland-Altman analysis, and Pearson correlation. DLC-derived measurements demonstrated strong agreement with manual scoring across all assays. ICC values were 0.929 (95% CI 0.648-0.971) for righting reflex, 0.965 (0.888-0.989) for negative geotaxis, and 0.958 (0.876-0.985) for wire hang. An automated behavioral analysis framework integrating DLC-based pose estimation with rule based quantification and supervised machine learning offers a reliable and objective alternative to manual scoring in neonatal HI models, enabling more efficient and reproducible behavioral assessment.

## Introduction

Hypoxic-ischemic (HI) brain injury in newborns is one of the leading causes of lifelong neurological disability, most notably cerebral palsy (CP) and global developmental delay, representing some of the most severe and prevalent disabilities in childhood [1–3]. Preclinical rodent models of neonatal HI are widely used to investigate the mechanisms underlying motor and cognitive deficits and to evaluate potential neuroprotective interventions [4–7].

Despite their central role in translational research, conventional behavioral assessments in rodent models present several well-recognized limitations. Many commonly used tests rely on subjective scoring systems that are prone to inter- and intra-rater variability, require extensive manual annotation, and are time-intensive to perform [8, 9]. In addition, quantitative analysis of motor behavior often depends on proprietary hardware and commercial software platforms with fixed experimental configurations and limited flexibility [10]. Such constraints can restrict experimental design, as investigators may need to adapt protocols to accommodate predefined equipment specifications and analysis pipelines [10]. When commercial tools fail to capture task-specific behaviors like subtle kinematic differences in neonatal rodent motor assays, researchers often resort to manual scoring, which is impractical for high-throughput analysis of continuous variables such as movement trajectories, velocity, and angular kinematics [11].

These limitations are particularly pronounced in neonatal HI models, where behavioral outcomes are often subtle, developmentally dynamic, and therefore difficult to quantify consistently [10, 12]. While established behavioral assays provide valuable measures of early motor function in neonatal rodents, their conventional evaluation often relies on manual timing or observer defined endpoints. As a result, measurement of these assays can remain vulnerable to subjective judgment, reduced temporal precision, and variability across raters and studies. These challenges highlight the need for quantitative, high resolution approaches that improve the accuracy, consistency, and reproducibility of behavioral measurements in neonatal HI models [13, 14].

Recent advances in computer science and deep learning have enabled markerless pose estimation approaches that overcome many of these limitations [15–18]. DeepLabCut (DLC), an open-source framework based on transfer learning with deep neural networks, allows accurate tracking of user-defined body parts across diverse animal species and experimental contexts [16, 19–22]. Due to its functionality, flexibility, and open-source resources, DLC has been widely adopted in behavioral neuroscience to analyze large sets of behavioral data without the need for physical markers or specialized equipment [16, 23–29].

In the present study, we focus on three well-established developmental motor assays—righting reflex (P8), negative geotaxis (P14), and wire hang (P16)—as they provide sensitive, age-matched measures of neuromotor maturation that are routinely impaired in neonatal HI models and clinically correlate with later cerebral palsy outcomes [6, 13, 14, 30–35]. To address the methodological challenges of these neonatal behavioral assessments, we evaluated the reliability of DLC-based pose estimation combined with rule-based analysis and supervised machine learning for automated, reproducible quantification of behavioral metrics in rodent neonatal HI models.

## Materials and Methods

### Study design and animals

Timed-pregnant Wistar rats were obtained from Charles River Laboratories (Wilmington, MA, USA) on embryonic day (E) 15 or 16 and housed in a temperature-controlled facility under a 12 h light/dark cycle with ad libitum access to food and water. The day of birth was designated postnatal day 0 (P0). On P1, litters were randomly culled to ten pups and balanced to achieve approximately equal numbers of males and females, minimizing inter-litter variability. On P7, each pup received a unique number and was randomly assigned within each litter to either the Sham group or the hypoxic-ischemic + placebo group (HI+PL). This procedure ensured unbiased group assignment while maintaining masking of investigators throughout surgeries, behavioral testing, and analysis. Sex was determined for all subjects. All experimental procedures were approved by the Institutional Animal Care and Use Committees (IACUC) of Brown University and were conducted in accordance with the National Institutes of Health Guide for the Care and Use of Laboratory Animals.

### Neonatal model of hypoxia-ischemia in rat

HI brain injury was induced on P7 using the Rice-Vannucci method, as previously described [36]. Briefly, pups were anesthetized with 4% isoflurane for induction and maintained at 2% isoflurane during surgery. Body temperature was maintained at 36.0 °C using a heating pad. A midline cervical incision (0.5-1 cm) was made, and the right common carotid artery was isolated and double-ligated with 5-0 silk sutures (Ethicon, Raritan, NJ, USA) in the HI+PL group. Sham pups underwent an identical surgical procedure without carotid ligation.

The incision was closed, and pups were marked for identification using tail ink injections (Neo-9, Animal Identification & Marking Systems, Hornell, NY, USA). Total anesthesia time was approximately 10-15 min, with carotid ligation completed within 5-8 min. Following surgery, pups were returned to the dam for 1.5-3 h of recovery before hypoxic exposure.

HI+PL pups were then placed in a hypoxia chamber (Biospherix, Parish, NY, USA) containing 8% oxygen balanced with nitrogen for 90 to 120 min. One non-ligated sentinel pup per litter was instrumented with a rectal temperature probe (RET-4, Physitemp, Clifton, NJ, USA) connected to a digital thermometer (TH-5, Physitemp Instruments, USA). Rectal temperature was recorded every 10 min and maintained as close to 36.0 °C as possible throughout hypoxia, as rectal temperature closely reflects brain temperature during HI exposure [37, 38]. Sham pups were placed in an identical chamber and exposed to room air for 90 to 120 min.

### Behavioral testing

All behavioral testing was conducted by investigators blinded to the treatment group. Animals were transported to the testing room and allowed to acclimate for at least 15 min prior to testing. Righting reflex and negative geotaxis were recorded from a top-down view using a monochrome GigE camera (Noldus, Leesburg, VA, USA). For the wire hang test, videos were captured from a lateral view using a consumer-grade camera (iPhone SE 3^rd^ Gen, Apple Inc., Cupertino, CA, USA). DLC-based pose estimation, as described by Mathis et al. (2018), is not constrained by the type of recording device, provided that video quality is sufficient for accurate tracking [16] [20].

### Righting reflex

The righting reflex test was performed on P8 to assess early vestibular and neuromuscular coordination following HI injury [35]. The righting reflex test was conducted according to established procedures reported in the literature [14, 33]. Each pup was placed on a flat surface, held in a supine position with all four paws upright by the experimenter. Upon release, the latency to right itself to a prone position with all four paws contacting the surface was recorded in seconds. Pups were allowed to flip for a maximum of 30 seconds, and trials in which pups failed to achieve a prone position within 30 seconds were considered failed and trials were repeated.

### Negative geotaxis

Negative geotaxis was assessed on P14 to assess vestibular function, neuromuscular coordination, and neurological development following HI injury [32]. The negative geotaxis test was conducted according to established procedures reported in the literature [14, 32]. Each pup was placed head-downward at the center of a 45° inclined wire mesh (48.26 x 50.8 cm [*L x W*]). Once the pup was stably grasping the wire mesh and supporting its body weight, the experimenter released the animal. The latency required to rotate at least 135° and orient the head upward was recorded in seconds.

### Wire hang

The wire hang test was performed in moderate HI model P16 rats to assess muscle strength, endurance, and motor coordination following HI injury [31, 34]. The wire hang test was conducted according to established procedures reported in the literature [14, 34]. Each rat was placed on a flat wire mesh, which was slowly flipped 180° by the experimenter. The flipped mesh was placed on top of a clear cage (30.5 x 30.5 x 18.4 cm [*L x W x H*]) lined with soft padding to prevent injury from the fall. Latency to fall was recorded in seconds.

### DeepLabCut

Markerless pose estimation was performed using DeepLabCut (DLC; version 3.0.0.rc13; Mathis Group, EPFL, Switzerland) following the protocols provided by the developers [16]. For each behavioral assay, 20 to 30 frames were sampled across time points and behavioral states to capture variability in posture, lighting, and occlusion. Frames were manually annotated for predefined body points relevant to neonatal posture and locomotion (**Fig. 1A**). Training parameters, including network architecture, number of training iterations, and batch size, were recorded, and model performance was evaluated using the built-in DLC metrics. Tracked x-y coordinates and likelihood values were exported as CSV files for downstream analysis using a customized open source pipeline based on DLCAnalyzer (https://github.com/ETHZ-INS/DLCAnalyzer)[22]. Behavioral measurements derived from DLC outputs were then compared with those obtained by manual human annotation (**Fig. 1B**).

**Figure 1.**
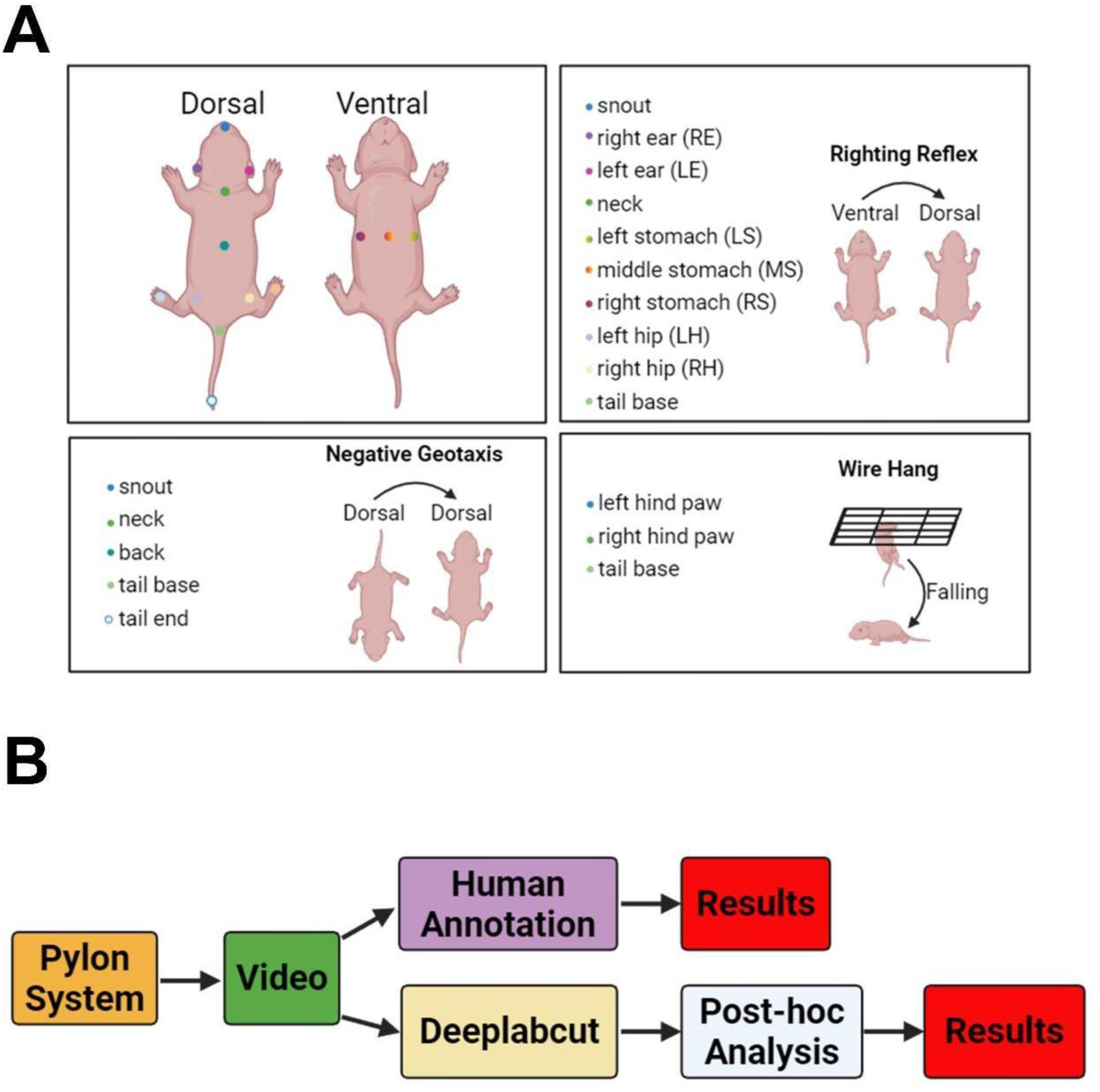
The labels used to train the DLC networks and behavioral analysis workflow. **A** Body points selected for DLC tracking across behavioral assays. The upper panel illustrates the full set of anatomical landmarks used for pose estimation from dorsal and ventral views, including the snout, right and left ears (RE, LE), neck, left, middle, and right stomach (LS, MS, RS), left and right hips (LH, RH), and tail base. Assay specific body points are shown for each behavioral test. For righting reflex, body points were used to quantify changes in body orientation during the transition from supine to prone. For negative geotaxis, body points were used to quantify body axis angle during turning. For wire hang, body points were used to quantify hanging and falling behavior. **B** Schematic overview of the behavioral analysis workflow. Video recordings were analyzed in parallel by manual human annotation and by DLC based pose estimation followed by post hoc computational analysis, and the resulting behavioral measurements were compared between methods.

### Coordinate-based measurement

DLC outputs consisted of frame-wise x-y coordinates and associated likelihood values for each tracked body part. Coordinates of each body part were processed using the DLCAnalyzer pipeline with minor modifications to accommodate study-specific behavioral definitions and measurements [22].

For negative geotaxis, five midline body points were tracked: snout, neck, back, tail base, and tail tip. To reduce the influence of low-confidence tracking from DLC, frames with a likelihood below 0.6 were interpolated. The resulting cleaned coordinate time series was used to compute a body axis vector. For each frame, the body axis vector from back to neck was calculated, and the angle between this vector and the positive vertical axis was computed. Completion of negative geotaxis was defined as the elapsed time from trial onset to the first “stable” arrival within a prespecified target range of 0-45 degrees relative to the positive vertical axis. Stability was defined as an angular velocity of the back-to-neck axis below 200 degrees per second sustained for at least 0.25 seconds. If the rat’s turn continued beyond the positive vertical axis, latency was instead recorded as the time to reach the vertical axis (180-degree turn). Turning latency was calculated by subtracting the human-measured trial onset from the computed completion time.

For wire hang, three body points were tracked: left hind paw, right hind paw, and tail base. To reduce the influence of low confidence tracking, body points with likelihood below 0.5 were excluded. Using DLC-derived coordinates, region-of-interest (ROI) occupancy was quantified for each tracked body part with frame-level temporal resolution. Trial onset was defined as the first frame in which any tracked body point entered or detected in the ROI. Trial completion was defined as the first frame in which no tracked body points remained within the ROI. Hanging duration was calculated as the time difference between trial onset and completion.

### Machine-learning based measurement

Wire hang and negative geotaxis behaviors were quantified directly from tracked body-part coordinates using rule-based definitions. In contrast, the righting reflex involves transitional postures that cannot be reliably captured using fixed thresholds on coordinate changes alone, so a supervised classifier was trained to predict righting-reflex state from pose-derived features based on the DLCAnalyzer pipeline with minor modifications [22].

For the righting reflex assay, ten body points were tracked: snout, right ear, left ear, neck, right stomach, middle stomach, left stomach, left hip, right hip, and tail base. To reduce the influence of low-confidence tracking, frames with likelihood values below 0.7 were interpolated, whereas frames with likelihood values below 0.2 were excluded. In addition, if more than three of the ten tracked body points had likelihood values below 0.2 at a given time point, the entire frame was excluded from analysis. The resulting cleaned coordinate time series was used for model training and inference. For each frame, acceleration was computed for each body point, and body-axis angles were calculated for three anatomical axes: tail base to neck, left stomach to right stomach, and left hip to right hip.

A separate dataset collected from previous righting reflex studies was used for machine-learning model development and included recordings from 84 rats housed across 14 cages. A neural network classifier was implemented using a Keras Sequential architecture consisting of two fully connected layers with 256 neurons in the first layer and 128 neurons in the second layer to predict righting-reflex state at the frame level. To incorporate short-term temporal context, predictions were generated using an integration period of 15 frames, corresponding to a 31-frame window centered on each frame. Each frame was classified into one of four mutually exclusive states: supine, transition, prone, or none. Frame-level labels obtained from manual scoring were used for training. The model was trained for 200 epochs with an internal 20% validation split, and the model with the highest validation accuracy was retained for all subsequent predictions.

### Statistical analysis

DLC-derived measurements were compared with manual scoring using three complementary approaches: Intraclass Correlation Coefficient (ICC), Bland-Altman analysis, and Pearson correlation coefficients. ICC (two-way, absolute agreement, single measures) was calculated using the irr package in R. Bland-Altman analysis was performed by computing the difference between manual and DLC-derived measurements, and 95% limits of agreement were reported. Pearson correlation coefficients were calculated to assess the linear relationship between automated and manual measurements. All other statistical analyses were performed using GraphPad Prism 10.5.0 (GraphPad Software Inc.). Righting reflex latency, negative geotaxis latency, and wire hang duration were compared between treatment groups using unpaired two-tailed t-test when data passed Shapiro-Wilk normality test; otherwise, the Mann-Whitney U test was performed. Statistical significance was defined as *p* < 0.05.

## Results

### Rat tracking

For each behavioral test, 241 frames for negative geotaxis, 238 frames for righting reflex, and 200 frames for wire hand, were manually labeled for DLC training using predefined body parts of interest from 10 to 20 rats. DLC training performances for each behavioral test were evaluated using root mean square error (RMSE), mean average precision (mAP), and mean average recall (mAR), which are summarized in **Table 1**.

### Righting reflex

The neural network classifier was trained on 84 righting reflex videos and achieved a highest validation accuracy of 0.77, indicating moderate frame level classification performance. **Fig. 2A** illustrates the temporal progression of behavioral states in a representative righting reflex video, comparing manually annotated frame level labels with the corresponding model predictions. Overall frame level classification performance is summarized in **Table 2**.

**Figure 2.**
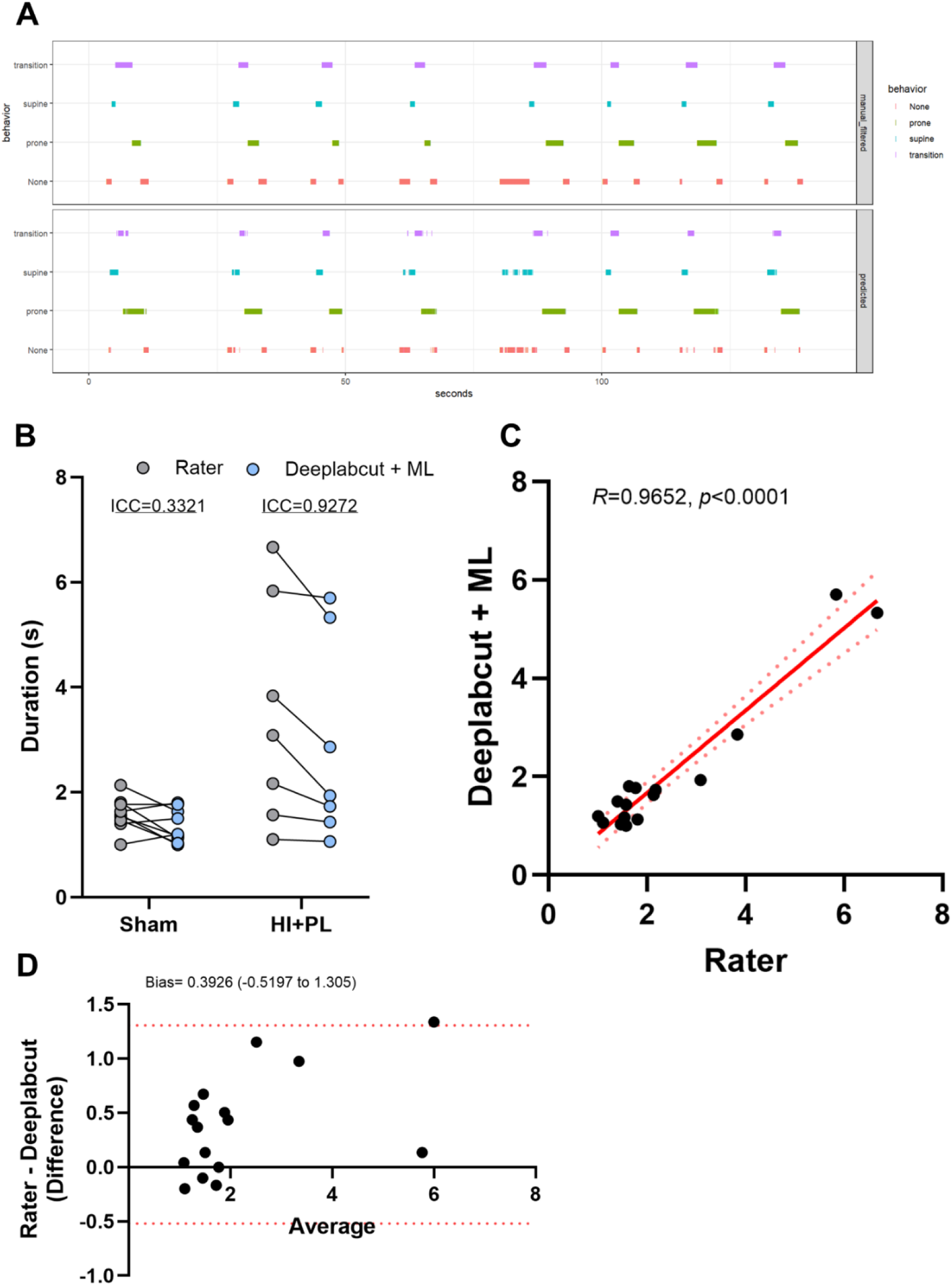
Deeplabcut + ML behavioral scoring performance on righting reflex. **A** Representative example of frame level behavioral state annotation over time across 8 trials, comparing manual labels and corresponding predictions from the DLC + ML pipeline. The top panel shows the manually annotated labels used as the ground truth for model training, whereas the bottom panel shows the corresponding frame level predictions generated by the DLC + ML model. **B** Comparison of righting reflex duration between manual human raters (gray circles) and the DLC + ML pipeline (blue circles) in the Sham and HI PL groups. ICC for each group are shown. **C** Pearson correlation analysis between manually measured righting latencies and those derived from the DLC + ML pipeline. A strong positive linear relationship was observed (r = 0.9652, *p* < 0.0001). **D** Bland Altman plot showing agreement between manual scoring and the automated DLC + ML pipeline. The y axis represents the difference between methods (manual minus DLC + ML), and the x axis represents the average of the two measurements.

Righting reflex performance was evaluated in 17 additional rats (9 Sham, 8 HI+PL) that were independent of the training dataset. Raw frame-level predictions were temporally smoothed using a hidden Markov model (HMM) to reduce frame-to-frame variability and penalize biologically implausible state transitions. The most probable sequence of behavioral states was then inferred using Viterbi decoding, a standard approach for optimal state sequence estimation in sequential models [39]. Righting latency was then computed from the post-processed state sequence as the time difference between the last supine frame and the first prone frame.

Manual measurements of righting latency were compared one-to-one with DLC-derived latencies. Paired comparisons demonstrated strong agreement between methods across individual animals, with high overall reliability (ICC = 0.9286, 95% CI 0.6479-0.9705). Correlation analysis further revealed a strong linear relationship between automated and manual scoring (Pearson’s R = 0.9652, 95% CI 0.9002-0.9881, **Fig. 2C)**.

Agreement between the two methods was further evaluated using Bland-Altman analysis (**Fig. 2D**), which showed a small mean bias of 0.3926 s, indicating that DLC measurements were slightly shorter on average than those obtained by the human rater. The 95% limits of agreement ranged from −0.5197 to 1.305 s, suggesting that most paired measurements differed by approximately ±1 s between methods. In subgroup analyses, agreement remained high in the HI+PL group (ICC = 0.9272, 95% CI 0.3331-0.9887, **Fig. 2B**), but was substantially lower in the Sham group (ICC = 0.3321, 95% CI−0.2046-0.7756, **Fig. 2B**). Both manual scoring and DLC-derived measurements detected significantly longer righting latencies in HI+PL animals compared with Sham controls (two-tailed t-test: Rater *p* = 0.0194; DLC *p* = 0.0331; Supplementary Table 1).

### Negative geotaxis

Negative geotaxis performance in 12 additional rats (6 Sham, 6 HI+PL) was measured manually by a trained human rater and compared one-to-one with DLC-derived latencies. **Fig. 3A** shows a representative example of negative geotaxis video analysis, in which the body axis angle relative to the vertical axis was tracked over time and used with a predefined threshold to determine task completion. Paired comparisons demonstrated strong agreement between methods across individual animals, with high reliability in duration measurements (overall ICC = 0.965, 95% CI 0.888-0.989; ICC = 0.9159, 0.5795-0.9854 for Sham; ICC = 0.9613, 0.7868-0.9934 for HI+PL, **Fig 3B**).

**Figure 3.**
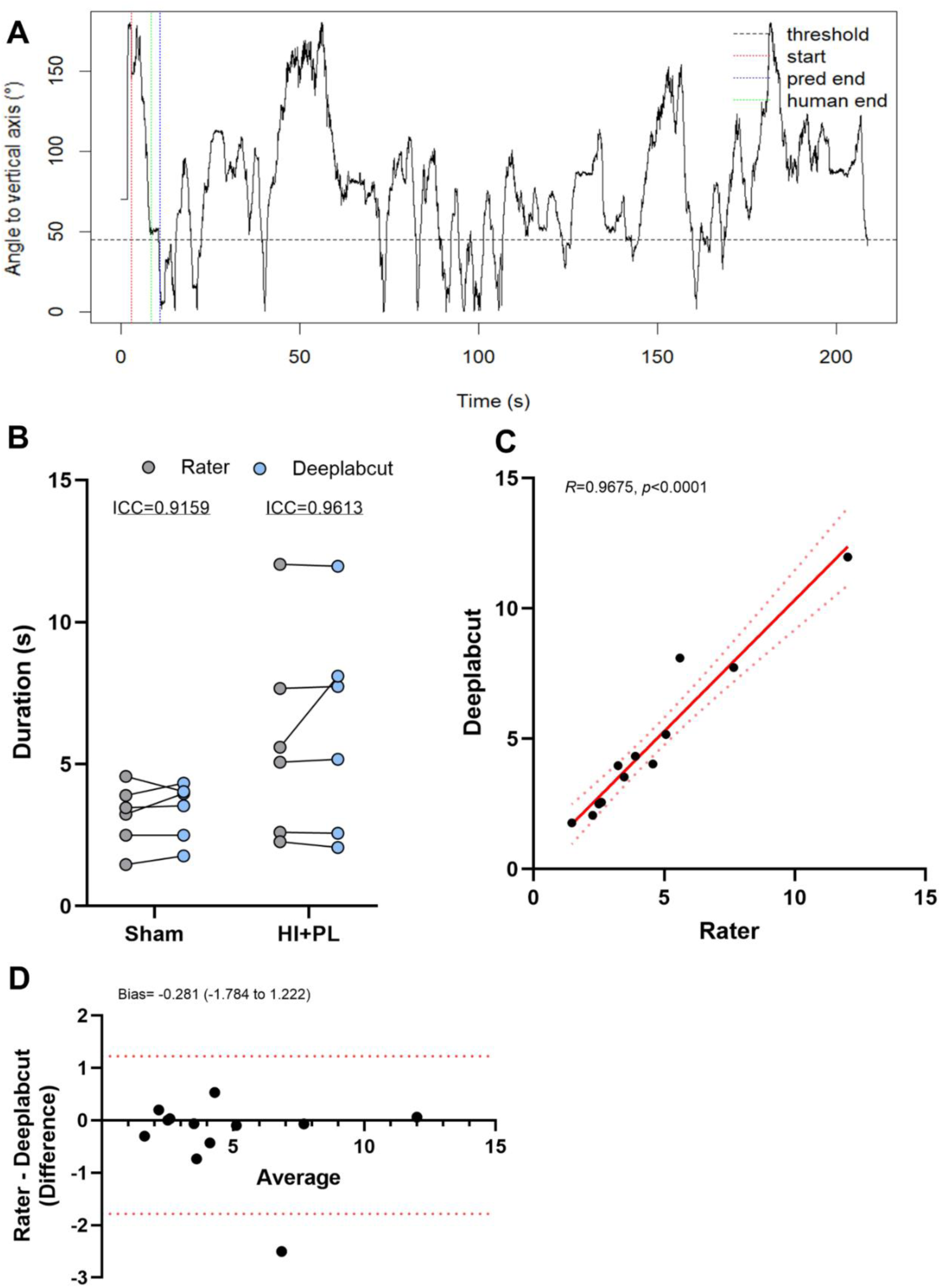
Deeplabcut behavioral scoring performance on negative geotaxis. **A** Representative example of body axis angle tracking during a negative geotaxis trial, illustrating the threshold based definition of task completion together with the predicted completion time and the manually annotated endpoint. **B** Comparison of negative geotaxis duration between manual human raters (gray circles) and the DLC pipeline (blue circles) in the Sham and HI PL groups. ICC for each group are shown. **C** Pearson correlation analysis between manually measured negative geotaxis latencies and those derived from the DLC pipeline. A strong positive linear relationship was observed (r = 0.9675, *p* < 0.0001). **D** Bland Altman plot showing agreement between manual scoring and the automated DLC pipeline. The y axis represents the difference between methods (manual minus DLC), and the x axis represents the average of the two measurements.

Correlation analysis further revealed an exceptionally strong linear relationship between DLC-based quantification and manual scoring (Pearson’s R = 0.968, 0.885-0.991, *p* < 0.0001; **Fig. 3C**).

Agreement between the two methods was further evaluated using Bland-Altman analysis (Fig. 3D), which demonstrated a small mean bias of -0.281 s, indicating that DLC measurements were slightly longer on average than those obtained by the human rater. The 95% limits of agreement ranged from -1.784 to 1.222s, suggesting that most paired measurements differed by less than approximately ±2 s between methods. No significant difference in latencies was observed between the Sham and HI+PL groups (Supplementary Table 1).

### Wire hang

Wire hang performance was evaluated in 14 additional rats (6 Sham, 8 HI+PL) independent from the training dataset. Latency to fall was measured manually by a trained human rater and compared one-to-one with DLC-derived measurements. The specific anatomical landmarks within a predefined ROI were tracked. **Fig. 4A** shows a representative example of wire hang video analysis, in which the left hind paw, right hind paw, and tail base were detected throughout the trial duration. Paired comparisons demonstrated strong agreement between methods across individual animals, with high reliability in duration measurements (overall ICC = 0.9583, 95% CI 0.8761-0.9849; ICC = 0.937, 0.6775-0.9899 for Sham; ICC = 0.967, 0.86-0.9929 for HI+PL; **Fig. 4B**).

**Figure 4.**
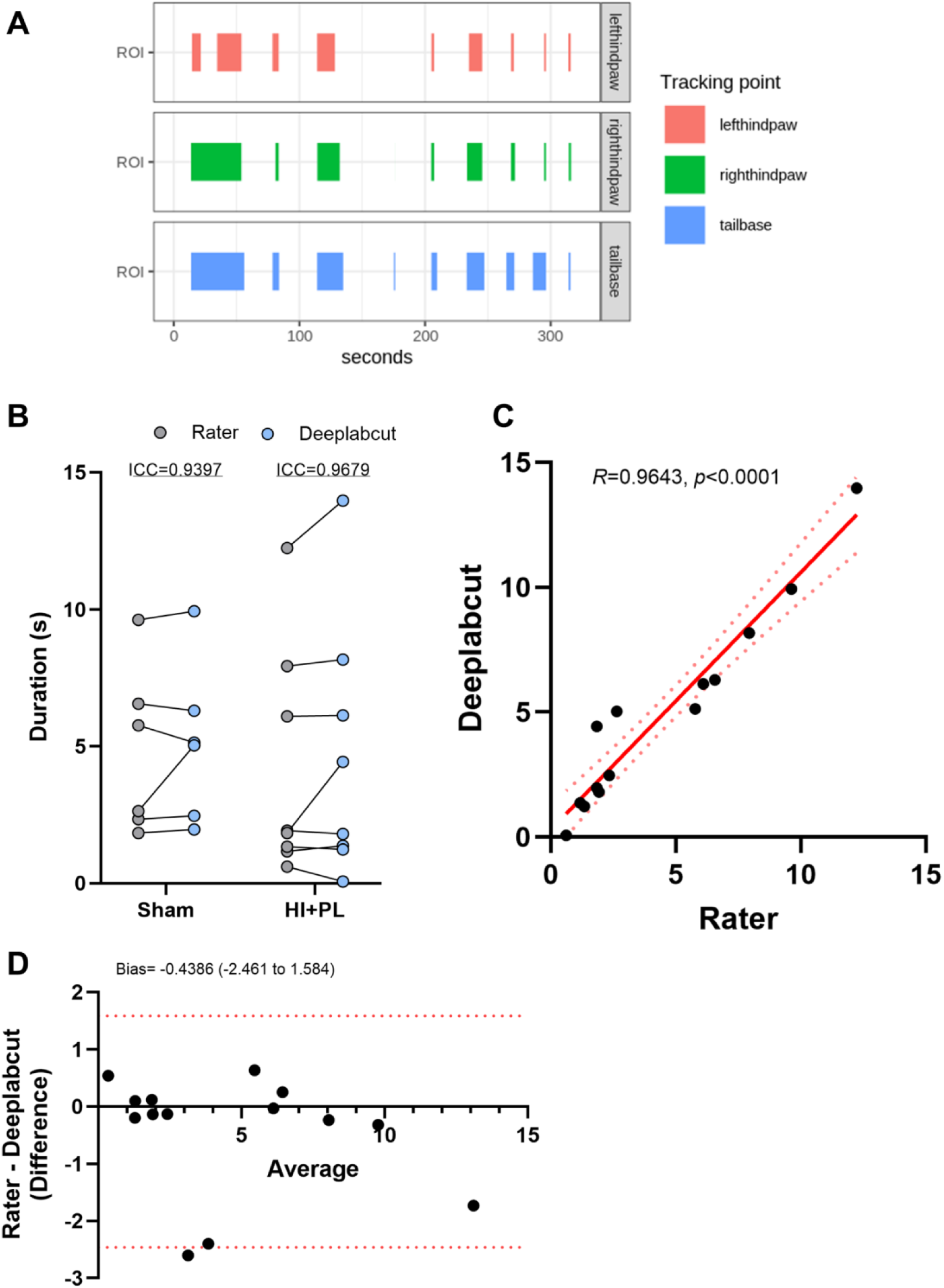
Deeplabcut behavioral scoring performance on wire hang. **A** Representative example of frame level detection of body points of interest during a wire hang trial. The left hind paw, right hind paw, and tail base were tracked over time to identify ROI occupancy duration during hanging behavior. **B** Comparison of wire hang duration between manual human raters (gray circles) and the DLC pipeline (blue circles) in the Sham and HI-PL groups. ICC for each group are shown. **C** Pearson correlation analysis between manually measured wire hang durations and those derived from the DLC pipeline. A strong positive linear relationship was observed (r = 0.9643, *p* < 0.0001). **D** Bland Altman plot showing agreement between manual scoring and the DLC pipeline. The y axis represents the difference between methods (manual minus DLC), and the x axis represents the average of the two measurements.

Correlation analysis further demonstrated a strong linear relationship between DLC-based quantification and manual scoring (Pearson’s R = 0.96, *p* < 0.01; **Fig. 4C**). Agreement between the two methods was further evaluated using Bland-Altman analysis (**Fig. 4D**), which demonstrated a small mean bias of −0.4386 s, indicating that DLC measurements were slightly longer on average than those obtained by the human rater. The 95% limits of agreement ranged from −2.461 to 1.584 s, suggesting that most paired measurements differed by approximately ±2 s between methods. No significant difference in latencies was observed between the Sham and HI+PL groups (Supplementary Table 1).

## Discussion

In this study, we developed and validated an open, customizable framework for automated behavioral quantification in a neonatal rat model with HI brain injury integrating markerless pose estimation with rule based and supervised learning approaches by using modified DLCAnalyzer pipeline[22]. Across three developmental tests, DLC-derived measurements demonstrated strong agreement with manual scoring, supporting the feasibility of reducing time intensive manual annotation while improving objectivity and reliability. A key advantage of this framework is its flexibility: rather than relying on proprietary systems with fixed configurations, the pipeline can be adapted to study specific definitions, camera views, and outcome measures using tracked key body points and transparent postprocessing rules.

For negative geotaxis and wire hang, behaviors were quantified using interpretable, coordinate-based definitions derived directly from tracked body points. Negative geotaxis turning latency, computed from the angle of the back-to-neck axis relative to positive vertical axis, showed excellent agreement with manual scoring and, in some cases, provided a more precise and internally consistent estimate than human observation. This advantage is most apparent when animals pause near, but do not cross, the fixed angular threshold. For example, with a 45° criterion, a pup may stop around 50° from vertical axis. Under these conditions, human raters may stop the timer based on the appearance of completion, even though the animal has not technically satisfied the predefined criterion. Such judgments rely on visual estimation and intuition, making them vulnerable to perceptual bias and difficult to apply consistently across trials and raters. By contrast, the automated approach applies the same geometric definition to every frame, reducing subjective judgment and improving reproducibility.

Additionally, the automated measurement of wire hang test by tracking body parts within a predefined ROI demonstrated strong agreement with human rater. Together, these results support the idea that many neonatal behavioral outcomes can be automated using transparent, rule-based definitions once high-quality tracking data is available, enabling reliable and standardized quantification across studies.

Unlike negative geotaxis and wire hang, which could be measured with simple rule-based definition, righting reflex poses a greater computational challenge because it involves rapid transitional postures, and brief intermediate states that are difficult to represent with a single coordinate threshold. To address this, we trained a supervised frame level classifier to label each frame as supine, transition, prone, or none. In our setting, raw frame level predictions were initially noisy and exhibited flickering, which is expected in multi-state prediction with a limited training dataset. We therefore incorporated temporal postprocessing using a hidden Markov model and Viterbi decoding to enforce sequential plausibility and reduce physically implausible state jumps.

At the outcome level, DLC-derived righting latencies showed high overall agreement with manual measurements in an independent validation cohort. Bland-Altman analysis demonstrated a small positive bias, suggesting modest underestimation of righting latency by the automated approach, while Pearson correlation indicated that the method preserved relative ordering across animals. Subgroup analyses revealed substantially lower agreement within the Sham group compared with the HI+PL group. This may reflect an inherently narrower range of latencies in Sham animals, in which short and tightly clustered durations make the measurements more sensitive to small timing differences. In addition, short-duration trials contain fewer frames in the transition state, which can further amplify sensitivity in agreement analyses.

Model performance may also have been influenced by characteristics of the training data. The dataset consisted of long recordings containing multiple consecutive trials from different pups, separated by inter-trial gaps. These intervals frequently included the experimenter’s hand, occlusions during repositioning, and occasional reattempts when a pup was inadvertently released before being fully positioned in a supine posture. Inclusion of such segments can increase label ambiguity, introduce non-task-related motion, and reduce the effective signal-to-noise ratio during training, particularly for boundary frames. These considerations suggest that automated righting reflex measurements may be most reliable when behavioral performance spans a broader range and highlight the importance of subgroup-specific validation when applying automated pipelines across experimental groups.

In addition to agreement analyses, the automated pipeline preserved biologically meaningful group differences. Manual scoring revealed significantly longer righting latencies in HI+PL compared with Sham animals (two-tailed t-test, *p* = 0.0194), and the same comparison using DLC-derived latencies remained significant (two-tailed t-test, *p* = 0.0331), indicating that the automated approach retained sensitivity to HI-related motor impairment at the group level.

Our findings align with prior work demonstrating that deep learning based behavioral analysis can approach human-level accuracy and in some cases outperform commercial tracking solutions, particularly when commercial tools fail to capture ethologically meaningful features or when experimental conditions deviate from standardized setups [22, 40–42]. Sturman et al. compared DLC based pipelines and supervised classifiers against human annotations and commercial platforms, showing that markerless pose estimation combined with machine learning can reach the accuracy of the human gold standard while reducing variability inherent to manual scoring [22]. Similarly, Rangoonwala et al. applied DLC to analyze motor behavior in parkinsonian rats, reporting ICC values of 0.9 and demonstrating that DLC-assisted scoring produced results comparable to manual measurements [42].

Despite these findings, this study has several limitations. First, validation was performed within a single experimental environment and imaging setup for each assay. Additional testing across laboratories, camera systems, lighting conditions, and experimenters will be necessary to establish the generalizability and robustness of the approach. Second, the righting reflex classifier achieved moderate-to-good frame-level accuracy (overall accuracy = 0.7694), and raw predictions exhibited frame-to-frame instability that required temporal postprocessing. Although hidden Markov model smoothing improves biological plausibility and reduces prediction flicker, future work should focus on improving the underlying classifier to minimize the need for extensive postprocessing. Potential strategies include expanding the size and diversity of the training dataset, improving labeling consistency for transitional states, engineering additional pose features that better capture body orientation and contact, and evaluating sequence-aware architectures that learn temporal structure directly.

Finally, the present study primarily evaluated whether automated measurements are comparable to conventional manual scoring. Future studies should focus on leveraging automated analysis to capture behavioral features that are difficult for humans to reliably quantify, thereby increasing sensitivity to detect subtle differences between groups and improving the prediction of neurodevelopmental outcomes following neonatal hypoxic-ischemic injury.

## Conclusion

In this study, we present a flexible and customizable framework for deep learning based behavioral analysis in a neonatal hypoxic ischemic brain injury model. By integrating DLC markerless pose estimation with coordinate based, rule driven quantification for negative geotaxis and wire hang, and supervised frame level classification for the more complex righting reflex, our approach enables automatic measurement of early locomotor and neuromuscular function. Across three different behavior tests, DLC-derived measurements showed generally strong agreement with manual scoring, with high ICC values and strong correlations, supporting that automated measurement can achieve near human level accuracy while reducing subjective judgment and rater dependent variability.

## Statements and Declarations

### Reproducibility and code availability

All pose estimation was performed using DeepLabCut (version 3.0.0.rc13). Downstream feature extraction and behavioral quantification were implemented using the open source DLCAnalyzer pipeline (https://github.com/ETHZ-INS/DLCAnalyzer) with study specific modifications. The machine learning workflow for righting reflex classification, including preprocessing, feature generation, model training and evaluation, and hidden Markov model postprocessing with Viterbi decoding, was implemented in R.

### Data availability

The datasets used and analyzed during the current study are available from the corresponding author upon reasonable request.

### Authors’ roles

B.L. was responsible for the statistical analysis and participated in drafting the manuscript; B.L and H.X performed data collection and analysis; B.L and H.X conducted AI model analysis and interpretation; B.L., H.X., B.W. M.L and X.C. participated in revising the manuscript; X.C. were responsible for the study design and data interpretation and finalized the manuscript. All authors read and approved the final manuscript.

