## Supplementary material for "Deep learning-based behavioral analysis in a neonatal rat model of hypoxic-ischemic brain injury": Tables

Table 1. Evaluation of DLC Training Model Performance Across Behavioral Tests.

| **Test** | **RMSE** | **mAP** | **mAR** |
| --- | --- | --- | --- |
| Righting Reflex | 5.73 | 82.57 | 89.17 |
| Negative geotaxis | 7.19 | 90.51 | 91.67 |
| Wire hang | 29.32 | 35.61 | 53.97 |

Table 2. Frame level classification performance (training set)

| **Class** | **Accuracy** | **Precision** | **Recall** | **F1 score** | **case (n)** |
| --- | --- | --- | --- | --- | --- |
| None | 0.7694 | 0.8721 | 0.6865 | 0.7682 | 10,362 |
| prone |  | 0.6091 | 0.9008 | 0.7268 | 6,842 |
| supine |  | 0.7596 | 0.8303 | 0.7934 | 3,612 |
| transition |  | 0.9163 | 0.7324 | 0.8141 | 7,007 |
