## Supplementary table for "Deep learning-based behavioral analysis in a neonatal rat model of hypoxic-ischemic brain injury"

Supplementary Table 1. P-values for comparisons between treatment groups across behavioral tests using manual scoring and DLC-based analysis.

| **Test** | **Rater** | **DeepLabCut** |
| --- | --- | --- |
| Righting Reflex | 0.0194 | 0.0331 |
| Negative geotaxis | 0.1129 | 0.0966 |
| Wire hang | 0.3626 | 0.4136 |

Normality was assessed using the Shapiro–Wilk test. Unpaired two-tailed t-tests were used for normally distributed data; otherwise, the Mann–Whitney U test was applied.
